# Investigating manta ray collective movements via drone surveys

**DOI:** 10.1101/2022.07.29.501955

**Authors:** Robert J. Y. Perryman, Culum Brown, Nicolò Pasian, Ashley J. W. Ward, M. I. A. Kent

## Abstract

Detailed observational research on free-ranging species of marine megafauna is required to understand their behavioural ecology, including how groups respond to environmental and anthropogenic pressures. New technologies are opening up potential for research on these species in the wild, especially on group-based and collective behaviours. Reef manta rays (*Mobula alfredi*) are socially interactive elasmobranchs that form groups in coastal reef habitats. Collective behaviours are likely important to their fitness, but may be disturbed by humans. Using small, remotely-piloted drones, we performed aerial observations of manta ray groups in Raja Ampat, West Papua. We empirically quantified patterns of collective movement including relative spatial positions, alignment, speed and leadership positions of conspecifics. We found unique patterns of spatial positioning, alignment and leadership, including differences between sexes, and high levels of local attraction, which were suggestive of distinct collective behaviour states. We suggest that ‘rules of interaction’ in manta rays vary at the individual level and can shift depending on local environmental and social conditions. Leader-follower behaviour likely has broad importance to cohesive movement and social behaviour in *M. alfredi*. We suggest that further studies on manta ray movement should consider utilising models of collective motion that capture group-level behavioural processes.

## Introduction

Understanding the behaviour of animals requires detailed observational research (Dawkins et al. 2007). Behaviour within groups reflects both density-dependent social forces, and how individuals balance costs and benefits of interacting with conspecifics (Stephens 2008; Farine et al. 2014; MacGregor et al. 2020). Individuals constantly assess their social environment to make optimal behavioural decisions that minimize predation risk, increase their foraging efficiency, and optimize their access to potential mates and suitable habitats (Miller et al. 2013; Marras et al 2015). Decisions on how to interact with groupmates are often made according to simple, self-organising rules (e.g. attraction to and alignment with others, positive and negative feedback, response thresholds) (Sumpter 2006; Herbert-Read et al. 2011; Silk et al. 2014). These ‘rules of interaction’ can drive complex collective behaviours, allowing groups to solve problems beyond the capacity of individuals (Ioannou 2017a), and enabling individuals to realise the fitness benefits of group membership (Couzin et al. 2005; Sumpter 2005; Lukeman et al. 2010). It is well recognised that high levels of alignment, reduced neighbour distances and cohesive movement enable transfer of information in groups of fish and promote efficient foraging and navigation (Ward et al. 2011; Herbert-Read et al. 2011). For example, in schooling fish, alignment in orientation and speed enables the rapid propagation of information gathered over a wide spatial range, without direct costs to individuals in directly sampling their environment (Couzin & Krause 2003; Ioannou et al. 2011; Berdahl et al. 2018). This improves collective navigation by dampening responses to small fluctuations in conditions.

Couzin et al. (2002) defined four collective movement states (‘swarm-like’, ‘torus’, ‘dynamic parallel swimming’ and ‘highly parallel directional alignment’) in schooling fish groups, based on levels of directional polarization and angular momentum. These states can occur interchangeably due to flexibility in rules of interaction, especially the size of ‘zones of attraction’ and ‘orientation’ that individuals adopt. Changes in collective behaviour enable individuals within groups to maximise fitness as local conditions change (Schaerf et al. 2017; Kent 2019), for example by increasing responsiveness to predators (Herbert-Read et al. 2017; Ioannou et al. 2017b). In homogenous groups, individuals with similar motivations usually respond similarly to their environment (Conradt and Roper 2005). The inherent complexity of many social systems, however, stems largely from differences between individuals (Jolles et al. 2020; MacGregor et al. 2020) that drives heterogeneity in responses to local conditions. Individuals with weaker social tendencies can exert ‘pulling power’ on others (Aplin et al. 2014; Brown & Irving 2014), while those with stronger motivation to be social can provide cohesion and promote stability in group structuring (Aplin et al. 2014; Ioannou et al. 2015). A range of emergent hierarchical and group-level effects (Herbert-Read et al. 2013; Jolles et al. 2020), including leader-follower polymorphisms (Aplin et al. 2014; Webster 2016), within-group assortment (Krause et al. 2000) and fission-fusion dynamics (Sueur et al. 2011; Silk et al. 2014) are common in social species living in variable environments. These may shift as a function of group size, cohesiveness and demographic composition (Sueur et al. 2011; Jolles et al. 2020). Leadership tendencies may be consistent and repeatable within individuals and reinforced by positive social feedback (Harcourt et al. 2009). In elasmobranchs, leadership during courtship, foraging and navigation (Jacoby et al. 2016; Stevens et al. 2018, Gore et al. 2019) can be driven by sex differences. Highly social species with complex cognitive abilities often exhibit more flexibility in their collective behaviour because individuals are able to recognise differences that exist between themselves and their groupmates and use this information to guide their decisions (Couzin et al. 2005; Ward and Webster 2016).

Quantitative analysis of leadership and collective behaviour is crucial to understanding group movement, information transfer, decision-making, exploration and predator evasion in social species (Couzin et al. 2005; King et al. 2009; Strandburg-Peshkin et al. 2015; Webster 2016; Jolles et al. 2020). Various mechanisms and functions of collective behaviours are well understood in captive fish under experimental conditions (e.g. Katz et al. 2011; Ward et al. 2011; Tunstrøm et al. 2013; Schaerf et al. 2017; Herbert-Read et al. 2017; Ioannou et al. 2017b; Kent 2019). Observational data on the behaviour of free-ranging marine species such as elasmobranchs, however, is notoriously difficult to collect. Fortunately, recent advances in remotely-piloted aircraft (RPA or hereafter ‘drone’) technology are allowing scientists to conduct detailed observations of marine animal behaviour *in situ*, without disturbance created by divers or in-water equipment (Anderson & Gaston 2013). Recently, the use of drones has advanced our understanding of shark (for review see Butcher et al. 2021) and ray (for review see Oleksyn et al. 2021) ecology, including presence (Hodgson et al. 2018), community structure (Kelaher et al. 2019) and population density (Kiszka et al. 2016; Tagliafico et al. 2020). Drones can be used as a tool to rapidly assess broad areas of potential elasmobranch habitat, monitor human-wildlife interactions (Colefax et al. 2019) and identify priority conservation areas (see Butcher et al. 2021 for review of drones in elasmobranch research). They can record high resolution tracks of focal animals (Raoult et al. 2018), and document inconspicuous and cryptic behaviours (e.g. mating, parturition) that are difficult to observe underwater (McCallister et al. 2020). Moreover, many elasmobranch species move in socially interactive formations (Sims et al. 2000; Guttridge et al. <u>2012</u>; Gallagher et al. 2014), and drones may provide excellent opportunities to study their social and collective behaviour (King et al. 2018). For example, Rieucau et al. (2018) used aerial images to document reef shark shoaling behaviour, while Gore et al. (2019) used aerial videos to document parallel swimming and close-following in basking sharks.

Reef manta rays (*Mobula alfredi*) are mobile pelagic rays that aggregate in surface waters of coastal regions during diel activities such as feeding, cleaning and courtship (Deakos 2010; Stevens 2016, Perryman et al. 2019). This makes them highly amenable to aerial monitoring but also vulnerable to anthropogenic impacts such as illegal fishing (Stewart et al. 2018) and unregulated dive tourism (Venables et al. 2016). Foraging, reproduction and predation in elasmobranchs often occur in groups, and may be easily disrupted by human activities (Roemer 2018). The Raja Ampat archipelago in West Papua, Indonesia is a critical habitat for a substantial reef manta population (Setyawan et al. 2020). The area is well protected from illegal fishing, but tourism has increased substantially in the past decade (Andradi-Brown et al. 2021). Reef mantas in Raja Ampat maintain social relationships that are linked to habitat preferences, sex-based assortment and fission-fusion dynamics (Perryman et al. 2019; Perryman et al. 2022). They frequent feeding sites and cleaning ‘stations’ which are focal points for social and courtship behaviours (Marshall & Bennett 2010; Stevens 2016). Courtship in manta rays involves male chasing of females through various stages (Stevens et al. 2018). Female *M. alfredi* in Raja Ampat appear to have stronger social preferences than males (Perryman et al. 2019), and collective behaviours may be influenced by this. Reef mantas appear to adopt several of the collective movement states defined by Couzin et al. (2002), e.g. parallel swimming and ‘cyclone feeding’ (Stevens 2016; Stevens et al. 2018). However, we do not yet understand how such group-based behaviours are characterised by the movement and spatial or hierarchical positioning of individuals, or whether collective states occur variably in response to local conditions. Research here may help to assess potential anthropogenic impacts on manta ray social behaviour.

Here we use drone-based video observations to: 1) assess the size and composition of groups of *M. alfredi* across multiple aggregation sites; 2) describe patterns of collective movement and 3) empirically quantify how behaviour at the individual level (movement speeds, alignment, relative positioning, leadership) differs in various group behavioural contexts. We investigate demographic and anthropogenic influences on responses to neighbour behaviour and emergent collective behaviour. We expected to find:

a. consistent patterns of relative positioning and alignment indicative of distinct collective behavioural states;
b. quantitative differences in movement speeds, group cohesion and leadership between these states;
c. individual and sex-based differences in movement speeds that are dependent on spatial and leadership positions;
d. stronger evidence for hierarchical leadership during ‘chasing’ behaviour than during other group-based behaviours;
e. that feeding behaviour is more common during running currents, and that feeding and ‘chasing’ behaviour is less common when groups were disturbed by humans.

## Methods

### Monitoring of manta rays via aerial surveys

Aerial surveys were performed on 80 different days between 22^nd^ January 2017 and 30^th^ May 2019 (between 08:00-18:00h), in the Dampier Strait area of Raja Ampat, West Papua (see Supplementary Information 1.1) according to a standardised protocol (see Supplementary Information 1.2). Surveys were conducted at known *M. alfredi* aggregation sites and designed to record the collective behaviour of groups of individuals. They were not performed during strong winds, rain, or in wave conditions higher than 1.25m, ensuring that manta rays were clearly visible to depths of ∼5-10m. Basic flight and wildlife observation data were collected from flight logs, and by visual inspection of recorded videos. For each survey we recorded the location, time of day, environmental conditions (including weather, sea state, and current conditions), using methods outlined in Perryman et al. (2019). We also recorded the number of rays of different sex and maturity status (see section 2.3), body size estimates (in pixels) using ImageJ (Rasband 1997-2002) (see Supplementary Information 1.3), the presence of boats (within ∼250m and ∼50m of manta rays) and the presence of human divers/snorkellers (within ∼50m and ∼10m of manta rays). In total we recorded usable data on groups of manta rays on 27 different days, at 5 different aggregation sites. See Supplementary Information 2.1 for results on group demographics.

### Tracking manta ray movements

We manually tracked the position and movement of all manta rays from 46 groups. We recorded a total of 5 hours, 58 minutes and 43 seconds of behavioural observations, but due to the necessity to occasionally reposition the drone when recording group-based movement, we analysed movement tracks from short clips (30-120 seconds) that were extracted from each ‘behavioural observation’ video, filmed at 24 frames/s. Data from 107 clips (range of 1-8 clips per video) was combined and considered as a single independent sample for each group. In all clips, the majority of individuals present at the aggregation site (counted during prior search) were clearly visible, and there was no movement of the drone or the frame of recording. We used VirtualDub 1.10 (Lee 2010) to convert clips into stacks of multiple image files (one per frame), scaled these to 1920 × 1080 pixels, and tracked movement using the ‘manual tracking’ plugin in Imagej using mouse autoclicker software (FastClicker; http://murgaa.com). We positioned the mouse cursor over the head of each individual ray, following its movement through the sequence of images to record an *(x, y)* coordinate on each click. We exported all coordinates and repeated the process for each individual ray in the group. Due to the variation in group sizes recorded, videos were recorded at different heights to fit all rays into the frame. These height differences had to be corrected to ensure that relative positions, alignment and speed could be compared between all observations. Therefore, following estimation of relative body sizes of all manta rays in each video, we scaled all coordinates according to the mean body length recorded for all individuals in that video. For further analyses, distances between rays are therefore recorded in ‘mean body lengths’, which is biologically appropriate where individuals vary in size.

### Recording individual characteristics

Demographic characteristics for all individual rays were recorded from observation of original video recordings, if possible. Distinct colour morph variants (‘melanistic’ and ‘typical’) (Venables et al. 2019) were recognised by their dorsal colouration (see Figure 1). Mature male rays were identified by the presence of calcified claspers, and mature female rays were identified by evidence from wing scars and pregnancies (as in Perryman et al. 2019). It was often difficult, however, to reliably identify the sex or maturity status of rays visually, so we used size comparisons to estimate these parameters (see Supplementary Information 1.3).

**Figure 1.**
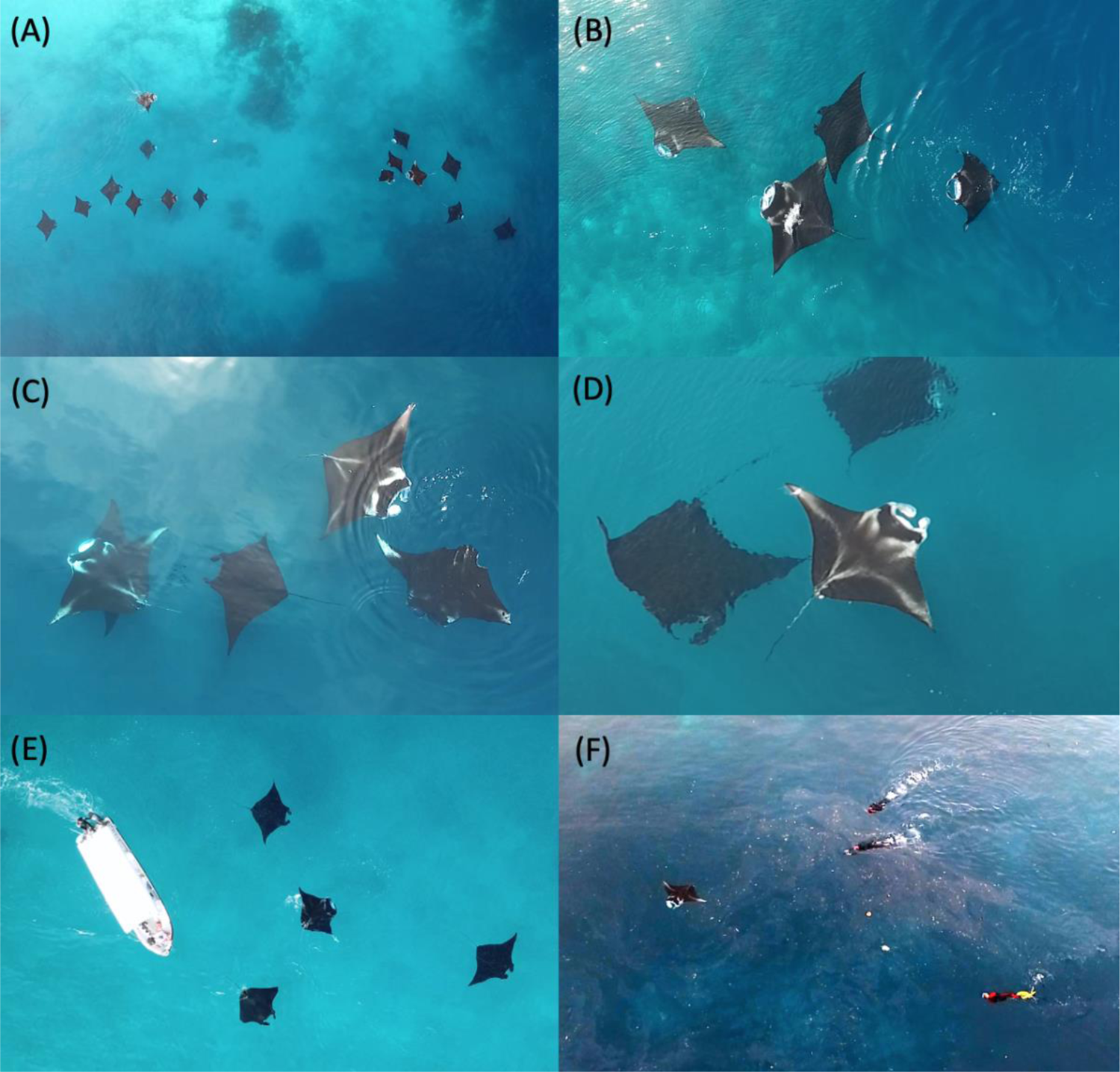
Images of manta rays captured during drone surveys. Panel A: image from typical altitude of behavioural observations (∼60m) showing division of larger group in two ‘subgroups’; Panel B: surface feeding behaviour (three typical morph, one melanistic morph manta rays); Panel C image from typical altitude for phenotypic data recording (∼15m), showing identification of female rays by wing scars; Panel D: identification of male typical morph manta ray by claspers; Panel E: disturbance of group of three melanistic morph and one typical morph manta rays by boat; Panel F: disturbance of manta ray by human snorkellers.

### Group behaviour types

To investigate potential differences between putative collective behaviours, we provisionally divided our recordings into three group ‘behaviour types’. These were defined based on movement characteristics of the majority of individuals visible, and included: ‘feeding’: where manta rays swam with cephalic lobes open, often with their upper jaw out of the water (Stevens 2016), either at a consistent heading angle or with repeated ∼180° turns (Stevens 2016); ‘chasing’: where manta rays led or followed behind another individual/s, emulating their speed and heading angle (similar to courtship initiation behaviour described by Stevens, 2016); or ‘swarm-like’ (hereafter ‘swarming’): where manta rays swam without any clear directional heading, and did not appear to interact socially except for slight directional changes when encountering conspecifics (Tunstrøm et al. 2013; Calovi et al. 2014). Categorisations were based on subjective interpretation and did not always capture the behaviour of all individuals, but highlighted differences that could be confirmed by subsequent quantitative analysis (see section 2.9).

### Patterns of movement and positioning within groups

To understand group-based movements and local rules of interaction, we recorded patterns of individual positioning and alignment as a function of relative partner position, and calculated basic measures of individual locomotion, group configuration, speed and relative alignment, using methods provided in sections S1.2, S1.3 and S1.5 of the supplementary material of Schaerf et al. (2017). We produced density heat plots from the perspective of a given focal individual by centring each ‘focal’ individual at the origin and rotating it so that its direction of travel was parallel to the positive *x*-axis, recording the positions of neighbouring rays relative to it, and repeating this for each frame and each individual in turn. These plots illustrated the probability of observing groupmates at particular (*x, y*) coordinates, relative to the focal individual. We used similar methods to create alignment heat plots showing both the mean relative directions of motion of partners at given (*x, y*) coordinates (illustrated with arrows) and a measure of the degree of focus of all the relative directions of motion observed for partners at given (*x, y*) coordinates (on a scale of 0, indicating high angular variance, to 1, indicating perfect alignment about the mean). We drew additional heat plots to show differences in chasing behaviour by sex, because initial results showed that individuals were most responsive to the position and alignment of others during this behaviour type, and because chasing behaviours are an important aspect of courtship in manta rays.

### Speed, directional alignment and leadership

We used coordinate data from behavioural tracks to calculate quantitative measures of speed and directional alignment as in Schaerf et al. (2017). The mean speed of all individuals in a group (hereafter ‘mean speed’) was taken as a measure of whole group movement. To calculate group-level polarization, we first smoothed all (x, y) coordinate data using a Savitsky-Golay filter (over 29 data points) in MS Excel. We recorded the arctangent (angle of direction of travel, in radians) for each individual over 5 frames and then calculated polarization in heading angle over all individuals (for each video frame), as the square-root of the summed sum-of-squares of the cosine and sine values for each individual arctangent value, divided by the number of individuals in the group, i.e. 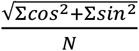. A value of 1 indicates that individuals were perfectly aligned, while a value of 0 indicates that individuals were completely out of alignment. We took the mean of polarization values over all frames as a measure of polarization for that group of rays. Individual-based values for alignment to the mean group heading were calculated as the numerical distance of the mean arctangent value for each individual over all frames, from the mean of means over all individuals. To investigate leadership, we used the R package *mFLICA* to infer dynamic following networks (Amornbunchornvej 2021) for each video. This technique used dynamic time warping on coordinate data from each video frame to detect leaders that initiate movement patterns, and followers of these patterns with a known time delay. After inferring following networks, we used the PageRank function in the R package *igraph* to calculate the relative importance of each individual as a leader (Amornbunchornvej & Berger-Wolf, 2018). A PageRank value of 1 indicates that an individual was followed by all other individuals throughout the recording, and a value of 0 indicates that it was not followed by any other individual. We used the network density of following networks as a measure of group-level coordination (a network density value of 1 indicates that all individuals followed the same pattern of movement, while a value of 0 indicates that all individuals moved completely independently).

### Regression-based analyses

We constructed various models in R (R Core Team 2019) to quantitatively explore how individual and collective movements were influenced by group demographics and by local social and environmental conditions, for all groups comprised of ≥4 individuals (smaller groups were excluded to focus on group-based collective behaviours). Details of model construction, testing and analysis are provided in Supplementary Information 1.4.

## Results

### Collective patterns of spatial positioning and alignment

There were qualitative differences in the relative positioning and alignment of manta rays within ‘chasing’, ‘feeding’ and ‘swarming’ groups. ‘Feeding’ and ‘swarming’ groups were more dispersed, though during feeding rays were likely to come very close together (within 0.5 BLs) (Figure 2a), with a strong mean focus of alignment in this small area, and some alignment behind and in front of a given focal ray (Figure 2d). In ‘swarming’ groups, rays were unlikely to be found within 2-3 BLs of a given focal individual (Figure 2b) and were not aligned in any direction (Figure 2e). During ‘chasing’, rays were frequently positioned approximately 0-3 BLs in front or behind, and within 1.5 BLs to the side of a given focal ray (Figure 2c) and in strong directional alignment (Figure 2f) from 0-5 BLs in front/behind, and 0-2 BLs to the side, focused in the areas of highest neighbour density. Female rays in ‘chasing’ groups were often positioned close together (within 2BLs, Figure 3a) and were well aligned directionally (Figure 3e). Male rays in ‘chasing’ groups were most likely to be positioned directly behind (at distances of 0-4 BLs), and not further than 2BLs to the side of a given focal female (Figure 3b) and were well aligned with females when directly behind and to the left side, but not well aligned on the right side (Figure 3f). They were also observed directly in front of females at a distance of 2BLs. Vice versa, female rays in ‘chasing’ groups were most likely to be positioned directly in front (rather than to the side) of a given focal male (Figure 3c), and aligned with males on the right side, and were also observed at 2BLs distance directly behind males. Males were more dispersed from each other at a range of distances in front or behind a given focal individual (Figure 3d) and were well aligned directionally but not when side-by-side. Patterns of collective behaviour observed here (tendency for males to follow females, and alignment of males on the left side of females) appear to be consistent with known ‘courtship initiation’ behaviour in reef manta rays, where a single (occasionally multiple) female/s will elicit chasing by multiple male rays (Stevens et al. 2018). Males appeared to approach and align with females from the left side (Figure 3f), which is consistent with studies showing behavioural lateralisation in mating behaviour-the vast majority of female wing scars from male bites during mating attempts are on the female’s left side (Marshall et al. 2010, RP pers. obs. in Raja Ampat).

**Figure 2.**
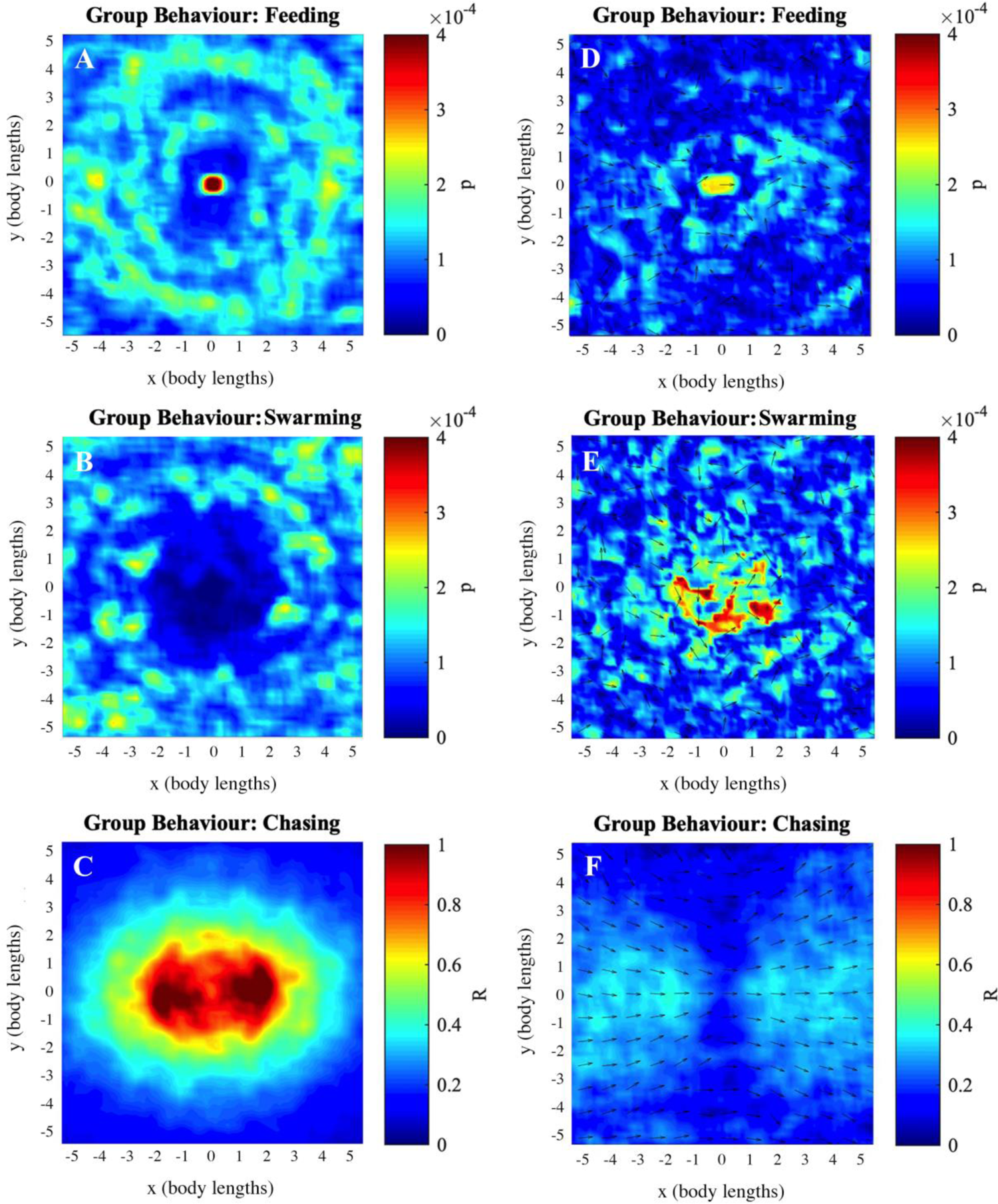
Relative positions and alignment of manta rays by collective behaviour type Panel A-C: relative frequency (estimated probability *p*) of partner manta rays at distances of 0-5 body lengths in any direction along a horizontal plane, relative to a given focal manta ray located at the origin and travelling parallel to the positive *x*-axis. Panel D-F: Mean relative direction of motion (arrows) and associated *R* values of partner manta rays based on their location relative to a given focal manta ray located at the origin and travelling parallel to the positive *x*-axis. *R* is a measure of the focus of all observed relative directions of motion within a particular observation, represented by the colour scale from 0 (greatest variance) to 1 (greatest focus about the mean). Panels compare groups performing different types of behaviour: feeding (Panels A, D), swarming (Panels B, E), chasing (Panels C, F). Warmer colours in heatplots denote higher relative frequencies of partner fish.

**Figure 3.**
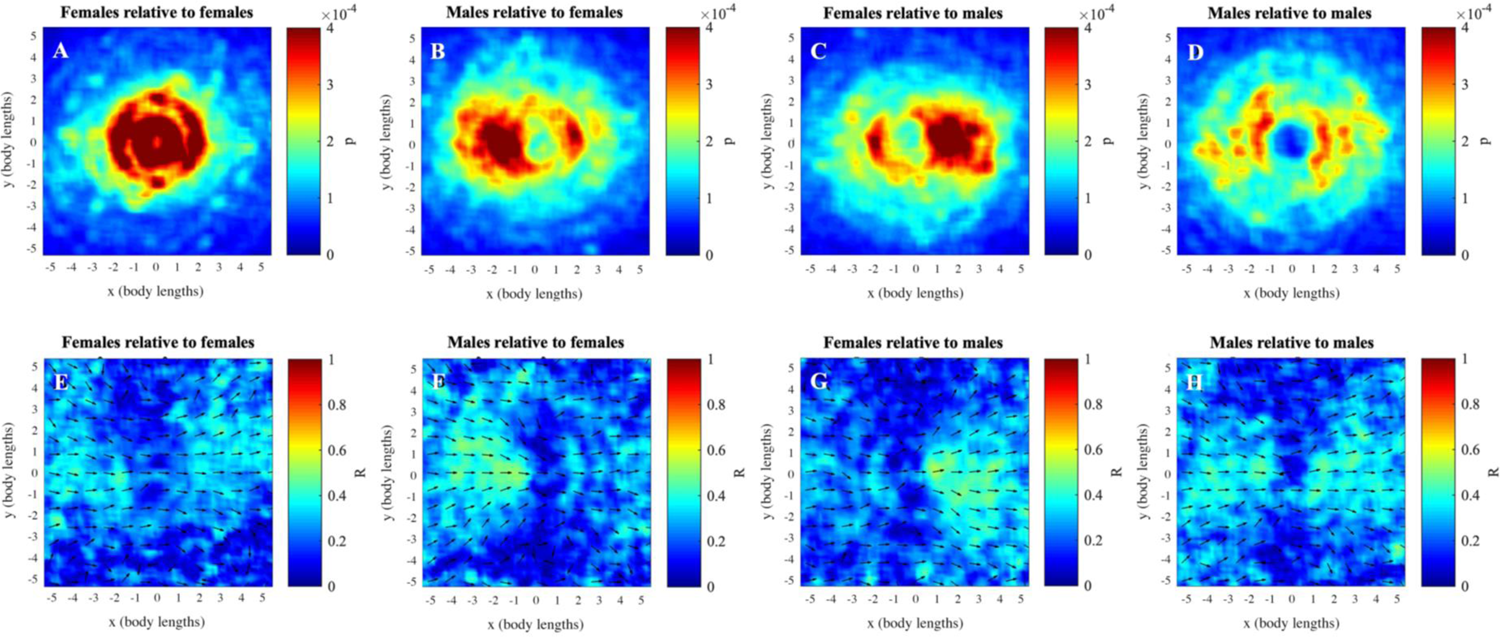
Heatplots of relative positions and alignment during ‘following’ behaviour Panel A-D: relative frequency (estimated probability *p*) of partner manta rays at distances of 0-5 body lengths in any direction along a horizontal plane, relative to a given focal mantaray located at the origin and travelling parallel to the positive *x*-axis. Panel E-G: Mean relative direction of motion (arrows) and associated *R* values of partner manta rays based on their location relative to a given focal manta ray located at the origin and travelling parallel to the positive *x*-axis. *R* is a measure of the focus of all observed relative directions of motion within a particular observation, represented by the colour scale from 0 (greatest variance) to 1 (greatest focus about the mean). Panels compare: females to a given focal female (Panels A, E), males to a given focal female (Panels B, F), females to a given focal male (Panels C, G), and males to a given focal male (Panels D, H). Warmer colours in heatplots denote higher relative frequencies of partner fish.

### Quantitative differences in collective behaviour

Results of multinomial models that quantified differences between group behaviour ‘types’ are shown in Table 1. Overall, results confirmed differences in collective alignment and leadership between the qualitatively classified described group behaviour ‘types’. During ‘chasing’, groups were larger (mean group size = 7.8 ± 2.96 individuals, 95% CIs), and more polarized (mean group polarization = 0.613 ± 0.097) than during ‘feeding’ or ‘swarming’. ‘Feeding’ groups were similar in size (mean 6.27 ± 2.9 individuals) than ‘swarming’ groups (mean 6.13 ± 1.25 individuals), with similar levels of directional polarization (mean feeding group polarization = 0.566 ± 0.133; mean swarming group polarization = 0.498 ± 0.063). Variability in leadership scores was similar for ‘chasing’ groups (SD of Pagerank values adjusted for group size = 0.401 ± 0.126) and ‘feeding’ groups (0.416 ± 0.163), and higher in ‘swarming’ groups (0.534 ± 0.109). Without controlling for group size, ‘feeding’ groups (SD of Pagerank values = 0.11 ± 0.06) had similar levels of variability in leadership scores to ‘swarming’ groups (0.092 ± 0.017), and both had higher variability than ‘chasing’ groups (0.067 ± 0.033). Results also suggested that external conditions influenced the emergence of different collective behaviours. ‘Feeding’ behaviour was most likely to occur, and ‘chasing’ behaviour least likely to occur, when currents were running. ‘Swarming’ behaviour was most likely to occur, and ‘chasing’ behaviour least likely to occur, when humans or boats were within 100m of the group of rays.

**Table 1.**
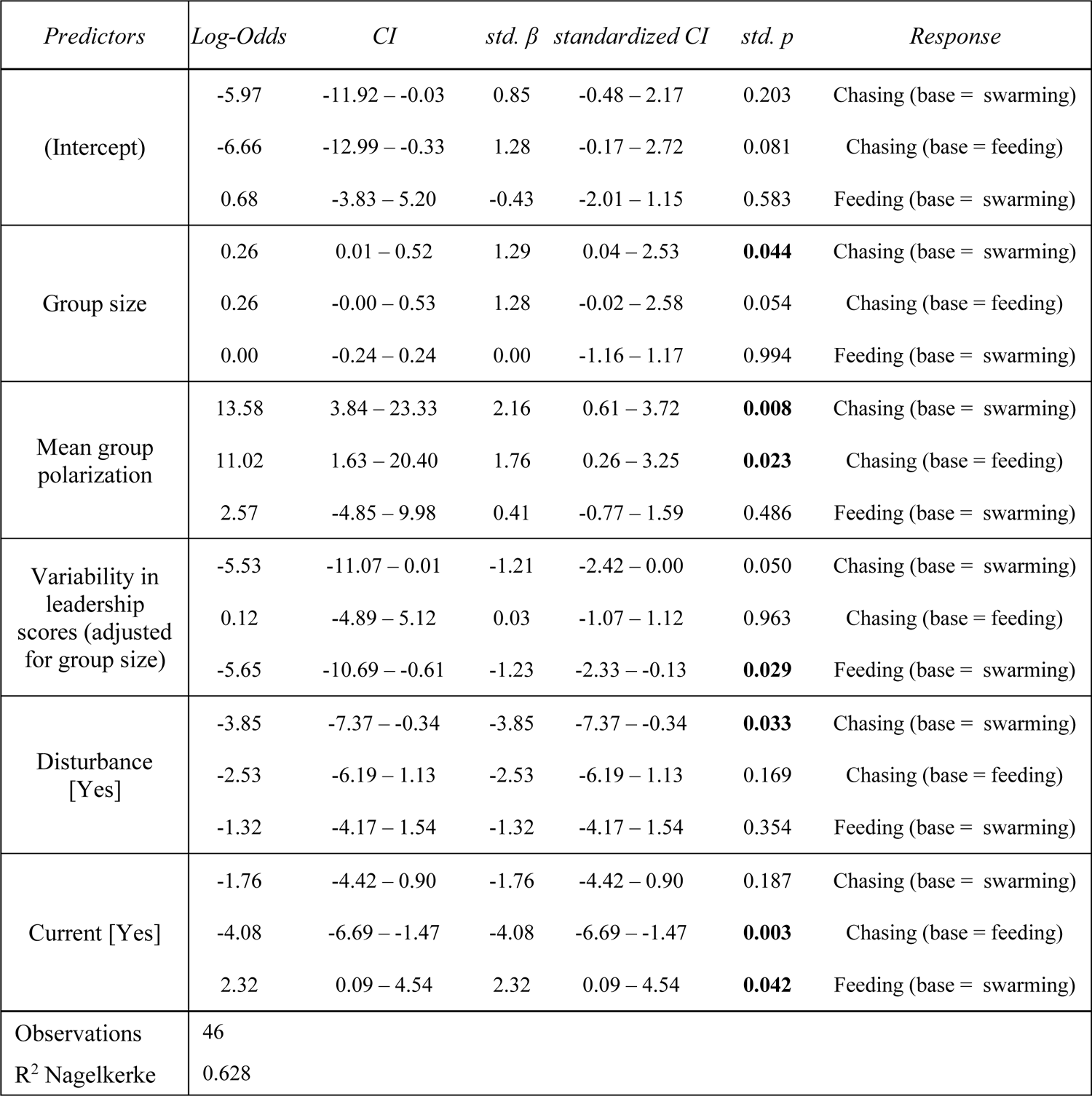
Quantitative differences in group properties by group behaviour classification

| *Predictors* | *Log-Odds* | *CI* | *std. $\beta$* | *standardized CI* | *std. p* | *Response* |
| --- | --- | --- | --- | --- | --- | --- |
| (Intercept) | -5.97 | -11.92 – -0.03 | 0.85 | -0.48 – 2.17 | 0.203 | Chasing (base = swarming) |
|  | -6.66 | -12.99 – -0.33 | 1.28 | -0.17 – 2.72 | 0.081 | Chasing (base = feeding) |
|  | 0.68 | -3.83 – 5.20 | -0.43 | -2.01 – 1.15 | 0.583 | Feeding (base = swarming) |
| Group size | 0.26 | 0.01 – 0.52 | 1.29 | 0.04 – 2.53 | **0.044** | Chasing (base = swarming) |
|  | 0.26 | -0.00 – 0.53 | 1.28 | -0.02 – 2.58 | 0.054 | Chasing (base = feeding) |
|  | 0.00 | -0.24 – 0.24 | 0.00 | -1.16 – 1.17 | 0.994 | Feeding (base = swarming) |
| Mean group polarization | 13.58 | 3.84 – 23.33 | 2.16 | 0.61 – 3.72 | **0.008** | Chasing (base = swarming) |
|  | 11.02 | 1.63 – 20.40 | 1.76 | 0.26 – 3.25 | **0.023** | Chasing (base = feeding) |
|  | 2.57 | -4.85 – 9.98 | 0.41 | -0.77 – 1.59 | 0.486 | Feeding (base = swarming) |
| Variability in leadership scores (adjusted for group size) | -5.53 | -11.07 – 0.01 | -1.21 | -2.42 – 0.00 | 0.050 | Chasing (base = swarming) |
|  | 0.12 | -4.89 – 5.12 | 0.03 | -1.07 – 1.12 | 0.963 | Chasing (base = feeding) |
|  | -5.65 | -10.69 – -0.61 | -1.23 | -2.33 – -0.13 | **0.029** | Feeding (base = swarming) |
| Disturbance [Yes] | -3.85 | -7.37 – -0.34 | -3.85 | -7.37 – -0.34 | **0.033** | Chasing (base = swarming) |
|  | -2.53 | -6.19 – 1.13 | -2.53 | -6.19 – 1.13 | 0.169 | Chasing (base = feeding) |
|  | -1.32 | -4.17 – 1.54 | -1.32 | -4.17 – 1.54 | 0.354 | Feeding (base = swarming) |
| Current [Yes] | -1.76 | -4.42 – 0.90 | -1.76 | -4.42 – 0.90 | 0.187 | Chasing (base = swarming) |
|  | -4.08 | -6.69 – -1.47 | -4.08 | -6.69 – -1.47 | **0.003** | Chasing (base = feeding) |
|  | 2.32 | 0.09 – 4.54 | 2.32 | 0.09 – 4.54 | **0.042** | Feeding (base = swarming) |
| Observations | 46 |  |  |  |  |  |
| R<sup>2</sup> Nagelkerke | 0.628 |  |  |  |  |  |

### Influences on individual and collective swim speeds

Average swim speeds for both sexes were similar (mean speed, females: 23.9 pixels/s, males: 26.5 pixels/s) but males swam faster than females in relation to their body size (female mean speed: 0.391 BL/s), IQ range: 0.238-0.532, male mean speed: 0.532 BL/s, IQ range: 0.34-0.683. Results of GLMMs with individual swim speeds as the response variable are shown in Table 2. Sex was an important predictor of manta ray swim speeds during ‘swarming’ (SES = 0.22, P = 0.01), but not in ‘feeding’ or ‘chasing’ groups, despite males having much higher average speeds in BL/s than females when chasing (female mean 0.317 +-0.047 BL/s, 95% CI; male mean 0.514 +-0.064 BL/s, 95% CI). This was likely due to high between-group variance (‘chasing’ τ_00_ = 0.28), with some groups being especially active, and low within-group variance in speed (‘chasing’ ICC = 0.89) especially in groups containing large numbers of males. Fixed effects were much less important in explaining overall variance than the random effect of group ID during ‘chasing’ (marginal R^2^ = 0.128, conditional R^2^ = 0.903), compared to during ‘feeding’ (marginal R^2^ = 0.398, conditional R^2^ = 0.589) or ‘swarming’ (marginal R^2^ = 0.518, conditional R^2^ = 0.753). Body length was an important predictor of movement speeds (in BL/s) during ‘feeding’, with larger individuals swimming relatively slower (SES = −0.31, P = 0.047), suggesting that smaller individuals may need to increase swimming effort to keep up with larger individuals. Manta rays with higher leadership scores tended to swim faster, especially during ‘feeding’ (female SES = 0.15, P = 0.24; male SES= 0.24, P = 0.11), but male leaders swam slower during ‘chasing’ behaviour (SES = −0.1, P = 0.075). During all behaviours, manta rays increased their swim speeds when out of alignment with the group mean heading (all effect sizes for different behaviour types positive, overall female SES= 0.11, P = 0.001; overall male SES= 0.06, P = 0.038) (see Figure S1 for linear models showing differences between behaviour types).

**Table 2.** Influences on movement speeds at individual level

| *Predictors* | Overall |  |  |  | Feeding groups |  |  |  | Swarming groups |  |  |  | Following groups |  |  |  |
| --- | --- | --- | --- | --- | --- | --- | --- | --- | --- | --- | --- | --- | --- | --- | --- | --- |
|  | *Est.* | *CI* | *Std. $\beta$* | *Std. P* | *Est.* | *CI* | *Std. $\beta$* | *Std. P* | *Est.* | *CI* | *Std. $\beta$* | *Std. P* | *Est.* | *CI* | *Std. $\beta$* | *Std. P* |
| (Intercept) | -1.05 | -1.84 – -0.26 | -0.91 | **<0.001** | -0.56 | -2.69 – 1.56 | -0.78 | **<0.001** | -1.64 | -2.62 – -0.65 | -0.82 | **<0.001** | -0.24 | -1.90 – 1.42 | -1.08 | **<0.001** |
| Males | 0.77 | 0.13 – 1.42 | 0.10 | 0.073 | 0.09 | -1.65 – 1.82 | 0.01 | 0.961 | 1.54 | 0.43 – 2.66 | 0.22 | **0.011** | 0.73 | -0.59 – 2.04 | 0.06 | 0.521 |
| Body length | -0.00 | -0.01 – 0.01 | -0.10 | **0.012** | -0.01 | -0.04 – 0.02 | -0.31 | **0.047** | 0.01 | -0.00 – 0.03 | 0.06 | 0.314 | -0.02 | -0.04 – 0.01 | -0.09 | 0.085 |
| Alignment (females) | 0.51 | -0.14 – 1.17 | 0.11 | **0.001** | 0.28 | -1.33 – 1.88 | 0.07 | 0.558 | 0.40 | -0.88 – 1.69 | 0.09 | 0.113 | -1.92 | -4.41 – 0.57 | 0.11 | 0.118 |
| Alignment (males) | 0.41 | -0.15 – 0.98 | 0.06 | **0.038** | 0.48 | -0.91 – 1.87 | 0.15 | 0.076 | 0.32 | -0.75 – 1.39 | 0.04 | 0.289 | -1.87 | -3.92 – 0.19 | 0.12 | **0.001** |
| Body Length * Alignment | -0.01 | -0.02 – 0.01 | -0.02 | 0.343 | -0.00 | -0.03 – 0.02 | -0.01 | 0.871 | -0.00 | -0.03 – 0.02 | -0.03 | 0.664 | 0.04 | 0.00 – 0.08 | 0.08 | **0.039** |
| PR (females) | 3.80 | 0.06 – 7.53 | 0.03 | 0.440 | 8.33 | -4.77 – 21.43 | 0.15 | 0.243 | 1.46 | -2.78 – 5.70 | 0.08 | 0.096 | 5.08 | -2.95 – 13.11 | 0.01 | 0.788 |
| PR (males) | 2.96 | -0.04 – 5.97 | -0.05 | 0.236 | 9.32 | -1.47 – 20.11 | 0.24 | 0.113 | 0.28 | -3.18 – 3.74 | -0.03 | 0.515 | 3.80 | -2.68 – 10.28 | -0.10 | 0.075 |
| Body Length * PR | -0.06 | -0.12 – -0.01 | -0.06 | **0.033** | -0.13 | -0.34 – 0.08 | -0.15 | 0.222 | -0.01 | -0.08 – 0.06 | -0.01 | 0.735 | -0.09 | -0.22 – 0.04 | -0.05 | 0.174 |
| Males * BL | -0.01 | -0.02 – 0.00 | -0.09 | 0.071 | -0.01 | -0.03 – 0.02 | -0.07 | 0.654 | -0.02 | -0.04 – 0.00 | -0.19 | 0.059 | -0.01 | -0.03 – 0.01 | -0.06 | 0.432 |
| Random Effects |  |  |  |  |  |  |  |  |  |  |  |  |  |  |  |  |
| $\sigma^2$ | 0.05 | | | | 0.08 | | | | 0.03 | | | | 0.03 | | | |
| $\tau_{00}$ | 0.16<sub>Video</sub> | | | | 0.04<sub>Video</sub> | | | | 0.03<sub>Video</sub> | | | | 0.28<sub>Video</sub> | | | |
| ICC | 0.76 |  |  |  | 0.32 |  |  |  | 0.49 |  |  |  | 0.89 |  |  |  |
| N | 28<sub>Video</sub> |  |  |  | 5<sub>Video</sub> |  |  |  | 11<sub>Video</sub> |  |  |  | 12<sub>Video</sub> |  |  |  |
| Observations | 232 |  |  |  | 50 |  |  |  | 74 |  |  |  | 108 |  |  |  |
| Marg. R<sup>2</sup> / Cond. R<sup>2</sup> | 0.203 / 0.809 |  |  |  | 0.398 / 0.589 |  |  |  | 0.518 / 0.753 |  |  |  | 0.128 / 0.903 |  |  |  |

PR = PageRank value indicating tendency to act as leader; Est. = beta coefficient; $\beta$ = standardized beta effect size; CI = confidence intervals for beta coefficient; $\sigma^2$ = Random effect variance; $\tau_{00}$ = between-video variance ICC = Intra-class correlation; Marg. = Marginal; Cond. = Conditional.

Results of linear regression models with group-based mean speeds as the response variable are provided in Table 3. Average movement speeds (in BL/s) were higher in groups with male-dominated sex ratios (overall SES = 0.2, P = 0.001) but not affected by group size. Average movement speeds in ‘feeding’ groups (SES = −0.27, P = 0.128) and ‘chasing’ groups (SES = −0.52, P = 0.3) were reduced when groups were more polarised directionally, and were strongly increased for ‘chasing’ groups when these were more cohesive (SES = 0.59, P = 0.25) and when leadership was less distributed (SES = −0.44, P = 0.02) between individuals. In ‘feeding’ and ‘swarming’ groups, average movement speeds were not strongly affected by levels of group cohesiveness or variability in leadership. Disturbance from boats or humans had strong positive effect on average movement speeds in ‘feeding’ (SES = 0.71, P = 0.14) and ‘swarming’ (SES = 0.63, P = 0.001) groups, but no effect on ‘chasing’ groups. Current had little effect on average movement speeds, except in ‘chasing’ groups, which swam much faster (SES = 1.89, P = 0.024) when currents were running.

**Table 3.**
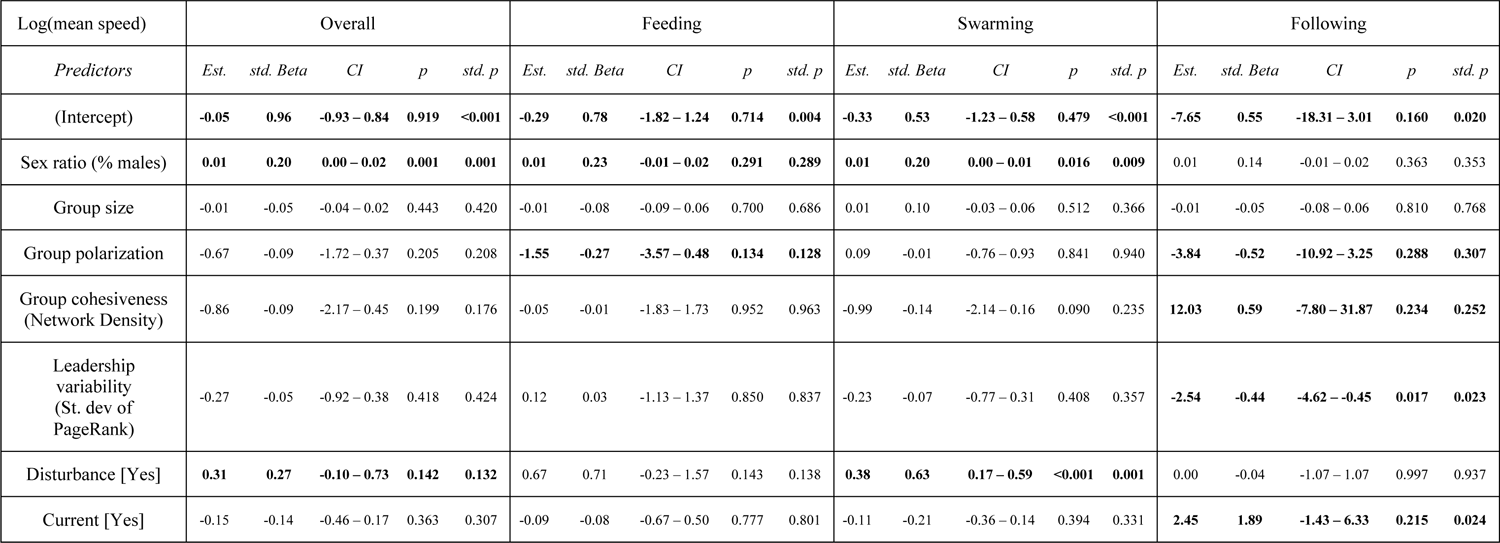
Influences on average movement speeds at group level

| Log(mean speed) | Overall |  |  |  |  | Feeding |  |  |  |  | Swarming |  |  |  |  | Following |  |  |  |  |
| --- | --- | --- | --- | --- | --- | --- | --- | --- | --- | --- | --- | --- | --- | --- | --- | --- | --- | --- | --- | --- |
| *Predictors* | *Est.* | *std. Beta* | *CI* | *p* | *std. p* | *Est.* | *std. Beta* | *CI* | *p* | *std. p* | *Est.* | *std. Beta* | *CI* | *p* | *std. p* | *Est.* | *std. Beta* | *CI* | *p* | *std. p* |
| (Intercept) | **-0.05** | **0.96** | **-0.93 – 0.84** | **0.919** | **<0.001** | **-0.29** | **0.78** | **-1.82 – 1.24** | **0.714** | **0.004** | **-0.33** | **0.53** | **-1.23 – 0.58** | **0.479** | **<0.001** | **-7.65** | **0.55** | **-18.31 – 3.01** | **0.160** | **0.020** |
| Sex ratio (% males) | **0.01** | **0.20** | **0.00 – 0.02** | **0.001** | **0.001** | **0.01** | **0.23** | **-0.01 – 0.02** | **0.291** | **0.289** | **0.01** | **0.20** | **0.00 – 0.01** | **0.016** | **0.009** | 0.01 | 0.14 | -0.01 – 0.02 | 0.363 | 0.353 |
| Group size | -0.01 | -0.05 | -0.04 – 0.02 | 0.443 | 0.420 | -0.01 | -0.08 | -0.09 – 0.06 | 0.700 | 0.686 | 0.01 | 0.10 | -0.03 – 0.06 | 0.512 | 0.366 | -0.01 | -0.05 | -0.08 – 0.06 | 0.810 | 0.768 |
| Group polarization | -0.67 | -0.09 | -1.72 – 0.37 | 0.205 | 0.208 | **-1.55** | **-0.27** | **-3.57 – 0.48** | **0.134** | **0.128** | 0.09 | -0.01 | -0.76 – 0.93 | 0.841 | 0.940 | **-3.84** | **-0.52** | **-10.92 – 3.25** | **0.288** | **0.307** |
| Group cohesiveness (Network Density) | -0.86 | -0.09 | -2.17 – 0.45 | 0.199 | 0.176 | -0.05 | -0.01 | -1.83 – 1.73 | 0.952 | 0.963 | -0.99 | -0.14 | -2.14 – 0.16 | 0.090 | 0.235 | **12.03** | **0.59** | **-7.80 – 31.87** | **0.234** | **0.252** |
| Leadership variability (St. dev of PageRank) | -0.27 | -0.05 | -0.92 – 0.38 | 0.418 | 0.424 | 0.12 | 0.03 | -1.13 – 1.37 | 0.850 | 0.837 | -0.23 | -0.07 | -0.77 – 0.31 | 0.408 | 0.357 | **-2.54** | **-0.44** | **-4.62 – -0.45** | **0.017** | **0.023** |
| Disturbance [Yes] | **0.31** | **0.27** | **-0.10 – 0.73** | **0.142** | **0.132** | 0.67 | 0.71 | -0.23 – 1.57 | 0.143 | 0.138 | **0.38** | **0.63** | **0.17 – 0.59** | **<0.001** | **0.001** | 0.00 | -0.04 | -1.07 – 1.07 | 0.997 | 0.937 |
| Current [Yes] | -0.15 | -0.14 | -0.46 – 0.17 | 0.363 | 0.307 | -0.09 | -0.08 | -0.67 – 0.50 | 0.777 | 0.801 | -0.11 | -0.21 | -0.36 – 0.14 | 0.394 | 0.331 | **2.45** | **1.89** | **-1.43 – 6.33** | **0.215** | **0.024** |

## Discussion

Here we present the first study of group-based behaviour in free-ranging manta rays (*M. alfredi*) using drone surveys. We reveal consistent patterns of collective movement with high levels of local attraction between conspecifics, driven primarily by leader-follower behaviour. We found quantitative differences in levels of polarization and leadership, providing evidence for distinct collective behaviour states. Our research suggests that these states occur interchangeably as individuals make active decisions to aggregate and interact socially (Perryman et al. 2019, 2022). This is likely driven by local social and environmental conditions (e.g. variable current conditions, group size and anthropogenic disturbance), differences in motivation between individuals, and flexibility in rules of interaction (Herbert-Read et al. 2013, 2017; Jolles et al. 2017, 2020). Negative correlations between swim speeds and alignment among all individuals (with similar effect sizes for different group behaviour types) suggest a general behavioural response where manta rays actively increased swim speeds to join collective behaviours, and decreased speeds when interacting with conspecifics. However, average movement speeds were correlated with polarization, cohesiveness and leadership in different ways for ‘feeding’, ‘swarming’, and ‘chasing’ groups, suggesting that modulation of swim speed during social interactions drives heterogeneity in collective movement.

During ‘swarming’ and ‘feeding’ behaviour manta rays were relatively well dispersed and did not align consistently with neighbours. However, many groups had high levels of local attraction and maintained strong directional alignment typical of polarized collective schooling (Couzin et al. 2002). These groups were elongated at a local scale with higher neighbour density and strongest alignment immediately in front or behind a focal individual, resulting in leader-follower (‘chasing’) behaviour. Such behaviour has been qualitatively described previously in *M. alfredi* and is known to occur during foraging (i.e. ‘chain feeding’) and courtship (‘i.e. ‘precopulatory chasing’) (Marshall and Bennett 2010; Deakos 2010; Stevens 2016; Stevens et al. 2018). Lower variability in leadership scores suggest that leadership was more distributed among different individuals during ‘chasing’ and ‘feeding’, compared to ‘swarming’ behaviour. This is somewhat surprising given that courtship behaviour in manta rays typically involves following of a single or small number of females by multiple males, and ‘chain feeding’ also implies hierarchical leadership. Influences on average swim speeds, however, suggest that ‘chasing’ groups with higher levels of hierarchical leadership were especially active, cohesive, and directionally unpolarised. This likely indicates courtship behaviour where the process of male mate selection by females in ‘mating chains’ includes rapid changes of direction and speed.

Leader-follower behaviour, however, is likely not only related to courtship. Instead, it may have broader importance to cohesive movement and social behaviour (Perryman et al. 2019), as suggested in filter-feeding basking sharks (Gore et al. 2019). Smaller group sizes and evidence for hierarchical leadership in slower moving ‘swarming’ groups, suggest ‘swarm-like’ (Couzin et al. 2002) movement as a social behaviour that may act as a precursor to chain feeding or courtship. Similarly, ‘chasing’ groups with more distributed leadership may occur as different small groups of socially interacting rays coalesce, with hierarchical leadership emerging later. The high between-group variance in individual swim speeds for ‘chasing’ groups (and much higher conditional R^2^ than marginal R^2^), compared to ‘swarming’ and ‘group feeding’ suggests that variability in leader-follower behaviour was greater between observations rather than within groups, suggesting a variety of functions in different social situations. Similarly, inconsistent patterns of positioning and alignment during ‘group feeding’ suggest flexible foraging methods that likely depend on local zooplankton density and oceanographic conditions (Stevens 2016). The high observed probability of feeding in close proximity to conspecifics may indicate ‘piggy-back’ feeding whereas manta rays are likely to be more dispersed during ‘ram’ feeding.

Females swim speeds were much more similar to male swim speeds during ‘chasing’ than ‘swarming’, and we found differences between the sexes in leadership-speed correlations (e.g. male but not female leaders decreased swim speeds during ‘chasing’ behaviour, and female but not male leaders increased swim speeds during ‘swarming’ behaviour), which suggest that sex-based behavioural differences were important in driving transitions between collective behaviour states. More social females (Perryman et al. 2019) may prioritise schooling behaviour (Brown & Irving 2014), as individuals with stronger social connections are likely to be followed by groupmates (Bode et al. 2012). Though male following of female rays was most common, same-sex ‘chasing’ interactions also occurred frequently. Female rays tended to position themselves close to other females, and in stronger directional alignment than for male-male interactions, indicating that females were more likely to maintain social contact and coordinate movements. This could be for a variety of reasons such as to resist sexual harassment (Jacoby et al. 2010; Perryman et al. 2019), or conversely to attract larger numbers of males. Speed modulation appeared to be related to hierarchical leadership variably according to overall patterns of collective behaviour. For example, during ‘chasing’ behaviour, average swim speeds were increased when groups moved cohesively (when most or all individuals followed the same leader/s), but the opposite was true for ‘swarming’ behaviour (decreased average speed for cohesive groups). Despite a larger sample size, confidence intervals for alignment-speed correlations were largest for individuals in ‘chasing’ groups, and confidence intervals for leadership-speed correlations were higher in ‘chasing’ compared to ‘swarming’ groups. This suggests that individuals vary in their response to leader-follower behaviour, and in their propensity to follow leaders. Individuals may also alter their behaviour in response to variable local conditions (e.g. the presence of sexually receptive females or an abundant food source). Behavioural flexibility in social species can enable balancing of coordinated social and collective behaviour with goal-oriented information-gathering and exploration (Aplin et al. 2014; Brown and Irving 2014). This is required for effective leadership (Ioannou et al. 2015), and is likely to be a key factor driving fission-fusion dynamics of manta ray groups in variable environments. Indeed, patterns of collective movement appeared to be strongly influenced by local conditions. For example, ‘feeding’ behaviour was more likely to occur when currents were running, likely due to increased efficiency of feeding on zooplankton.

Our observations showed that various types of collective behaviour were common in the upper 5-10m of the water column in areas adjacent to cleaning stations. These should be priority areas for management interventions, particularly as large, socially interactive and coordinated groups of manta rays are likely to attract the attention of tourism operators. Increased boat traffic in areas of high manta ray activity is likely to increase vulnerability to boat strikes and propellor injuries, while large numbers of snorkellers or SCUBA divers may disturb socially interactive or collective behaviours that likely have important fitness-related functions in courtship and feeding. ‘Swarming’ behaviour was more likely to occur, and ‘chasing’ behaviour less likely to occur, when boats or divers were close to groups of rays. This could be due to less active ‘swarming’ groups being easier for tourism operators to access, but we also found that average swim speeds of manta rays in ‘feeding’ and ‘swarming’ groups were increased under anthropogenic disturbance. Results therefore suggest that manta rays actively respond to disturbance from humans, and that this facilitates changes in collective behaviour, perhaps dissipation of slow-moving leader-follower behaviour. In this context, further understanding of how, where and why collective behaviours occur, and the impact of human disturbance on them, is important.

In summary, collective movement appears to be an important aspect of the biology of manta rays that is likely to affect all aspects of their life histories, as well as being vital to foraging and courtship. Our research demonstrates the utility of drone surveys in understanding the collective behaviour of wild elasmobranchs. Further studies might use models of collective decision-making (e.g. Farine et al. 2014; Ioannou et al. 2017a) or information transfer (Ioannou et al. 2011) to explain movements and dispersal in response to environmental drivers and anthropogenic pressures. Frequency-dependent selection for leader-follower behaviour is known to be an important driver of personality and dominance-based relationships in animal groups (Aplin et al. 2014), and likely occurs in rays (Pini-Fitzsimmons et al. 2021). Research on between individual variation and within-individual behavioural flexibility in leadership is required to link individual and group-based behaviours, which should lead to better understanding of fission-fusion dynamics in manta ray groups. If possible, leadership models should attempt to incorporate repeated measures of behaviour on specific individuals (Aplin et al. 2014; Jolles et al. 2017), though the inability of drones to identify individual animals is a major limitation compared to in-water surveys. A solution here might be to combine aerial surveys with data from remote underwater cameras or animal tag telemetry. Drones may also be a useful tool in managing manta ray populations. Regular drone-based aerial surveys of priority areas could achieve multiple goals, including: better understanding of behaviour and human-wildlife interactions; monitoring and recording of boat traffic and diver behaviour; and identifying unsustainable or malign human activities. Better understanding of collective behaviour processes and human impacts should enable better conservation (Westley et al. 2018), including informed assessment of local vulnerability, and the development of effective rules and regulations for marine tourism operators.

## Notes

### Competing Interest Statement

The authors have declared no competing interest.

### Summary of Updates

Small changes to text and author list

