## Supplementary Information for "Investigating manta ray collective movements via drone surveys"

### 1. Methods

### 1.1. Location and sampling methods

Our study site in Raja Ampat, West Papua is a shallow-water marine habitat including areas of coral reef, seagrass beds and sandy bottom, among small islands and sandbanks. Manta rays form feeding aggregations and visit cleaning station sites here from November to April each year. Various tourism activities (boating, SCUBA diving, snorkelling), as well as local subsistence fishing occur throughout the area during daylight hours.

### 1.2. Drone survey protocol

We used a DJI Phantom III Advanced and a DJI Mavic Pro quadcopter drone (www.dji.com) to perform aerial surveys. Both drones were equipped with an onboard camera, GPS, compass and tri-axis gimbal, allowing for flights at consistent speed and altitude, and stable video recording at an adjustable angle. Cameras recorded 2706 x 1520 or 1920 × 1080 pixel wide-angle video at 24 frames/s, for the duration of all survey flights. Survey flights were launched from islands, sandbanks, built wooden structures or from a small speedboat. They began with a ‘search’ phase and progressed to ‘behavioural observation’ phase if a group of manta rays was detected. On reaching a known aggregation site, the drone was flown slowly over it at an altitude of 80-120m whilst filming at 90° angle to the water surface, for at least 30 seconds. This was sufficient to detect manta ray presence due to the shallow water habitat and high contrast of manta rays’ dorsal colouration against the substrate. If manta rays were observed, the drone was positioned hovering directly above the largest group of rays visible, at a height of 40-80m (lower if the group was small, higher if the group was large and dispersed), with the camera facing straight down (important to avoid parallax issues and to accurately record the relative size of each ray). The ‘behavioural observation’ phase then commenced, with ~8-10 minutes of continuous video recorded. Occasionally the movement of manta rays out of shot necessitated slight adjustment to drone position. Following this (if battery levels permitted), we lowered the altitude of the drone to approx. 20m, and hovered over each individual ray in turn to provide higher resolution imaging and more accurate data on sex, body size and colour morph. At this height the sound of the drone is indistinguishable from background underwater noise, and therefore marine animal behaviours are unlikely to be affected by it (Butcher et al. 2021).

### 1.3. Estimating maturity and sex through size-based comparisons

For individual manta rays whose sex or maturity status we could not determine visually by standard methods, we used size based comparisons to estimate this. We recorded and ranked the relative size of each individual, using this as a proxy for maturity. We used the ruler in ImageJ to measure body length (in pixels) from the edge of the upper jaw to the trailing edge of pelvic fins, in image frames where the body was at the water surface, close to the centre of the frame (this minimised distortion errors due to parallax or bending of the body surface). Body length (BL) is proportional to disc width (DW) in manta rays (Deakos 2010b) and was considered more reliable here because pectoral fins are bent during swimming, whereas the body is rarely flexed lengthways. Mature male *M. alfredi* in Raja Ampat range from 2.25-3.75m DW, mature females range from 2.5-4.5m DW, and immature males are not larger than 2.75m DW (Perryman et al. 2019). We therefore considered rays that both lacked visible claspers (i.e. immature males) and were >125% the size of the smallest identified mature male (i.e. larger than 2.75m DW) in each clip as mature females. Assuming that females reach sexual maturity at larger size than males, we also classified all rays that were <60% the size of the largest identified mature male (i.e. smaller than 2.25m DW), or <50% the size of the largest individual as immature. All remaining rays, and those for which body size could not be estimated (e.g. due to not being at the surface or centre of frame), were recorded as sex or maturity status ‘unknown’. It was not possible to identify distinct individual rays from the dorsal surface, so we could not determine whether individuals were recorded repeatedly between videos.

### 1.4. Regression models

Data exploration was carried out following the protocol described in Zuur, Ieno & Elphick (2010), and models were validated by checks of residual and other diagnostic plots (QQ plots, Cullen-Frey graphs) using the *fitdistrplus* package (Delignette-Muller & Dutang 2015), with the appropriate probability distribution for regression models chosen through AIC-based comparisons. We used the ‘dropterm’ function in the R package ‘MASS’ to test all combinations of fixed effect variables, and removed variables that increased AIC from final models. We calculated standardized effect sizes to judge the relative importance of fixed effects. To calculate quantitative differences in collective behaviour between the group ‘types’, we performed multinomial (logistic) regression using the R package ‘nnet’, with group behaviour type (‘feeding’, ‘swarming’ or ‘chasing’) as the response variable, and various group-level characteristics as predictor variables. These included the mean of individual speed values (BL/s) as a measure of group activity, group-level polarisation, the standard deviation of within-group PageRank values (weighted by group size), following network density, group size (number of individuals), sex ratio (% males), ‘disturbance’ (swimmers, diver or boat presence within 100m of the group), and current strength (none/slight or medium/strong). Variables ‘mean speed’, ‘sex ratio’ and ‘following network density’ increased AIC and were removed from the final model. Results compared group behaviour ‘types’ directly (i.e. ‘chasing’ is compared to ‘feeding’, not to values averaged over all behaviours).

Manta rays often make rapid changes to swim speed in response to external stimuli (Perryman et al. 2021; pers. obs.). During group interactions these are likely to reflect individual responses to conspecifics. Therefore, in an attempt to determine factors influencing rules of interaction, we constructed regression models with individual movement speed as the response variable. Individual speed values were non-normally distributed, and right-skewed, so we constructed Gamma generalized linear mixed models (GLMMs) with a log link function, using the ‘glmmTMB’ package (Brooks et al. 2017). We modelled each individual’s speed (BL/s) as a function of its sex, body length, alignment (deviation in radians from the mean group heading) and leadership position (PageRank value divided by group size). We included interactions between sex and alignment/leadership, and between body length and alignment/leadership, because we expected differences between mantas of difference sex/age in their movements in response to social stimuli. The variable ‘colour morph’ increased AIC and was removed from the final model. Because our observations consisted multiple individual manta rays within groups, we included the observation ID (Video #) as a random intercept to model the dependency structure among observations of individuals in the same group. We constructed separate models for individuals in ‘feeding’ (n_inds_ = 50), ‘swarming’ (n_inds_ = 74), and ‘chasing’ (n_inds_ = 108) groups, as well as an overall model including data on individuals (n_inds_ = 232) within all groups. To investigate potential influences on collective behaviour at the group level, we constructed Gaussian linear models with the natural log mean group speed in BL/s (of all individuals in each group) as the response variable (AIC comparison indicated best fit to the log-normal distribution). Predictor variables were group polarization, group size, group sex ratio, within-group variability in leadership (standard deviation of pagerank values), group cohesiveness (following network density), disturbance (swimmer, diver or boat presence within 100m of the group) and current strength (none/slight or medium/strong) as predictor variables. We constructed separate models for ‘feeding’ (n_groups_ = 5), ‘swarming’ (n_groups_ = 11), and ‘chasing’ (n_groups_ = 12) groups, as well as an overall model including data from all groups (n_groups_ = 28).

### 2. Results

### 2.1. Group demographics

We analysed movement tracks from a total 303 individual reef manta rays in 46 groups. Group sizes ranged from 2-21 individuals (mean 6.72). 117 individuals were recorded as females (38 from pregnancies or wing scars, 81 from size comparison), 106 as males (all from direct inspection of their claspers), and 80 as ‘unknown sex’. 195 individuals were recorded as mature adults, 13 as juveniles, and 95 as ‘unknown maturity’. 33 groups contained both females and males, with four being comprised only of females, three only of males, and six of unknown sex composition. Sex and maturity ratios were similar to those found in previous research in Raja Ampat using photo-identification methods (Perryman et al. 2019). Mature female reef manta rays were on average around 20-25% larger than mature males (IQ ranges, females: 60-66 pixels, males: 48.5-53.5 pixels), which was consistent with previous studies utilising in-water size estimation (Perryman et al. 2019), and drone-based size estimation utilising a floating reference measure (Setyawan et al. 2022). Seven of our observations had humans or boats within 50m of the group of rays. 30 surveys were in calm current conditions, and 16 were in moving current conditions (stronger than ~ 0.2m/s). For ‘feeding’ rays, we recorded tracks over 16,731 frames for 88 individuals in 17 groups, equivalent to 1 hour, 10 mins and 4 seconds. For ‘swarming’ rays, we recorded tracks over 22,190 frames, for 86 individuals in 15 groups, equivalent to 1 hour, 29 mins and 57 seconds. For ‘chasing’ rays, we recorded tracks over 43,184 frames, for 128 individuals in 15 groups, equivalent to 3 hours, 18 mins and 40 seconds. However for analysis of movement speeds (Tables 2 and 3) we removed groups containing fewer than four individuals, reducing the number of groups to 28 (5, 11 and 12 respectively for feeding, swarming and chasing groups).

2.1. Movement speed

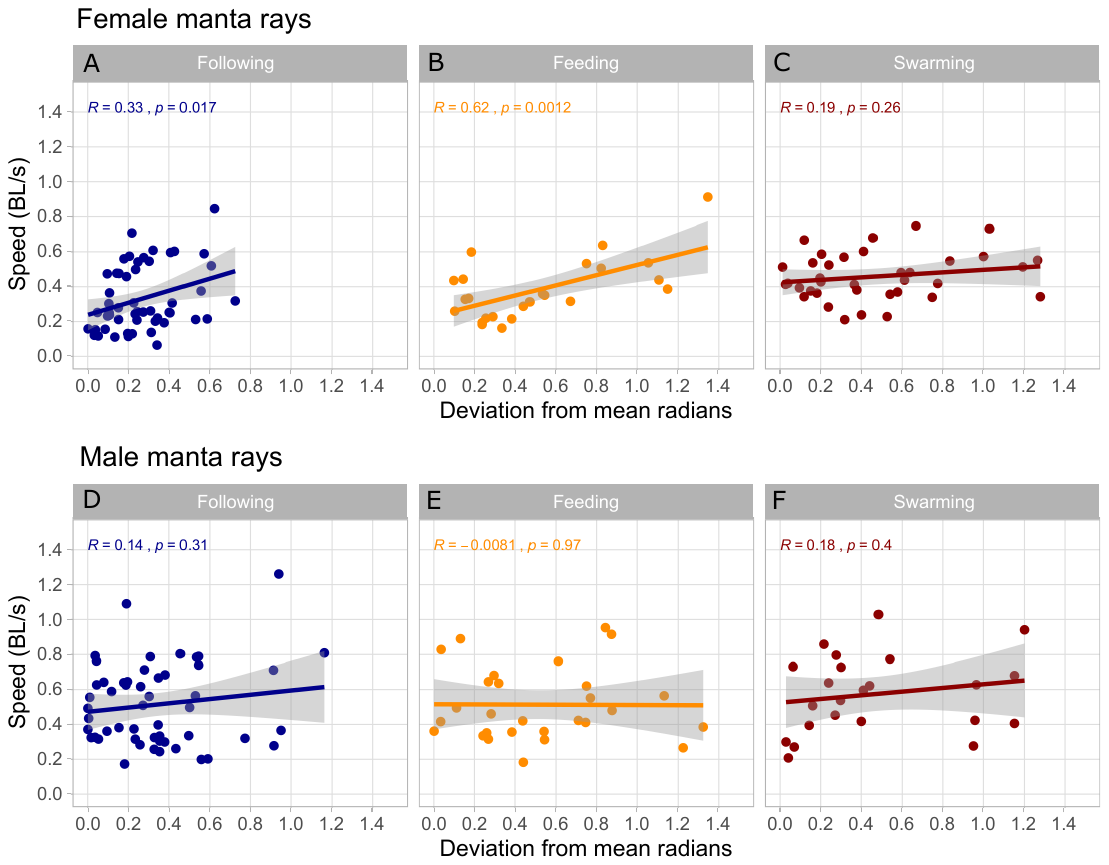

Figure S1. Linear models showing swim speeds during different behaviour types**.** Speeds in body lengths/s (average over whole observation) as a function of individual deviation from the mean direction of travel for all rays in the group (average over whole observation), for female (upper panels) and male (lower panels) reef manta rays during ‘following’ (Panels A, D), ‘feeding’ (Panels B, E) and ‘swarming’ (Panels C, F). Except for males in ‘feeding’ groups, both sexes increased speeds when out of alignment with group mean heading, and this was especially pronounced for females in ‘following’ and ‘feeding’ groups.
